# Correlations of neural predictability and information transfer in cortex and their relation to predictive coding

**DOI:** 10.1101/2025.05.23.655705

**Authors:** J. P. Fiorenza, L. Le Donne, C. Schmidt-Samoa, E.-C. Heide, C. von der Brelie, V. Malinova, N. K. Focke, M. Wilke, L. Melloni, C. M. Schwiedrzik, M. Wibral

**Author notes:** Contributed equally as co-first authors (J. P. Fiorenza, L. Le Donne).

## Abstract

Predictive-coding like theories agree in describing top-down communication through the cortical hierarchy as a transmission of predictions generated by internal models of the inputs. With respect to the bottom-up connections, however, these theories differ in the neural processing strategies suggested for updating the internal model. Some theories suggest a coding strategy where unpredictable inputs, i.e., those not captured by the internal model, are passed on through the cortical hierarchy, whereas others claim that the predictable part of the inputs is passed on. Here, we addressed which neural coding strategy is employed in cortico-cortical connections using an information-theoretic approach. Our framework allows for quantifying two core aspects of both strategies, namely, predictability of inputs and information transfer, through local active information storage and local transfer entropy, respectively. A previous study on the neural processing of retinal ganglion cells connected to the lateral geniculate nucleus showed a coding for predictable information, captured by an increase in the information transfer with the predictability of inputs. Here, we further investigate predictive coding strategies at the cortical level. In particular, we analyzed LFP activity obtained from intracranial EEG recordings in humans and spike recordings from mouse cortex. We detected cortico-cortical connections with increasing information transfer with the predictability of inputs in recorded channels from frontal, parietal and temporal areas in human cortex. In the mouse visual system, we detected connections exhibiting both an increase and decrease in the information transfer with input predictability, although the former was predominant. Our evidence supports the presence of both predictive coding strategies at the cortical level, with a potential predominance of encoding for predictable information.

**Summary:** The ability of the brain to infer the hidden causes of sensory experiences has been conceptualized within the computational framework of predictive coding. This framework explains perceptual inference and learning as a process of constantly updating an internal model of the world. Predictive coding describes cortical activity as a communication of sensory evidence and predictions generated from prior expectations. While different views of predictive coding agree on the communication of prior expectations throughout cortex, they differ in how internal expectations are updated. One view states that, to update the internal model, the cortex propagates the mismatch between the expected neural activity and the actual neural response to sensory stimuli. In contrast, another view suggests that the cortex propagates the match between the expected neural activity and the actual neural response. In this work, we were able to tease apart these two views, both in human cortex and the mouse visual system, using information theory. We observed that the brain predominantly propagates expected information, i.e., the match between prior expectations and incoming sensory inputs.

## 1 Introduction

The “predictive principle” [1] states that the brain performs perceptual inference and learning by adjusting an internal generative model of the world which captures the statistical regularities of the environment. This principle assigns a purpose to the observed hierarchical organization of the cortex [2] by assigning different roles to the feedforward and feedback connections [3]. The feedforward connections, i.e. those from lower to higher cortical areas, convey the sensory information obtained from the environment. Whereas the feedback connections, i.e. those from higher to lower cortical areas, communicate predictions made by the generative model of the world. This generative model encodes at each cortical level a representation that aims to predict the neural activity at the level below.

The feedforward communication of sensory evidence and feedback transmission of the predictions of the internal generative model were formally described within the computational framework of predictive coding [4, 5].

One specific flavor of predictive coding conceptualizes each cortical level as composed of two types of units, named error and representation units (e.g. [4, 3]). The representation units signal predictions via feedback connections about the neural activity of the cortical level below. The error units compute the mismatch between these predictions and the neural activity received from lower hierarchical levels, producing prediction errors that are transmitted further via feedforward connections.

We will refer to the feedforward communication of prediction errors as the predictive processing strategy of error-coding.

However, feedforward connections could also convey neural activity patterns that are in agreement with the predictions of the generative model [6]. In this case, the feedback connections also transmit the internal predictions which are compared with the incoming sensory data. However, this comparison enhances neural activity patterns that represent sensory information that is predictable in the given context. This enhancement in the neural response to sensory features that match our internal predictions has been described by the biased competition model [7] and Adaptive Resonance Theory [6]. We will refer to the feedforward communication of predictable patterns of neural activity as the predictive processing strategy of reliability-coding.

These two coding strategies differ fundamentally in what is passed on from lower to higher cortical areas, but both rely on the principle of top-down predictions. The error-coding strategy implies a transfer of unpredictable information, whereas the reliability-coding strategy implies a transfer of predictable information.

Thus, at the center of these strategies there are two core concepts, i.e., information transfer and predictability.

In this work, we quantified information transfer and predictability using two information-theoretic measures from the Local Information Dynamics Framework [8]. Information transfer was quantified using Transfer Entropy (TE) [9] and predictability was measured using Active Information Storage (AIS), and their respective localized variants (LTE, LAIS) [8]). AIS was computed on a neuronal source (i.e. a neuron or a brain area) and TE was computed from a neuronal source to a target (a neuron or a brain area). This framework allows us to move beyond some of the previous limitations in the study of predictive processing. For example, in the study of the neural mechanisms involved in predictive processing, the experimental design usually requires the control of the stimulus-predictability. This limits the study of predictive processing to only those datasets collected while controlling the stimulus-predictability. In contrast, the Local Information Dynamics Framework allows us to investigate predictive processing without the control of the stimulus-predictability by studying the internal neural predictability. We note that this internal predictability will ultimately be also the only predictability available to neurons or circuits.

In this study, we directly quantified predictability from neural activity (given that throughout the brain, neuronal inputs come from other neurons) thus adopting a neuro-centric perspective. Moreover, by quantifying information transfer and predictability directly from neural activity, we were able to expand predictive processing research to experimental designs where the stimulus-predictability is not controlled.

The two coding strategies detailed above can be teased apart by their information-theoretic footprint in the following way: the error-coding strategy implies a decrease in the information transfer with increasing neural-predictability and it is represented in a negative correlation between TE and AIS; the reliability-coding strategy implies an enhancement of information transfer with increasing neural-predictability and it is represented in positive correlation between TE and AIS. This relationship between information transfer and neural-predictability was previously defined as Storage-Transfer Correlations (STC) and provides a signature of predictive processing [10]. Evidence of positive STCs were first detected between the retina and the LGN in the cat visual system, providing a signature of a reliability-coding strategy at the sub-cortical level [10].

In this work, we move forward by providing evidence for the presence of STCs at the *cortical* level in the human brain and mouse visual system. We specifically asked: can significant STCs be detected at the cortical level? Do cortico-cortical connections follow an error-coding or reliability-coding strategy?

We show that STCs can be detected in cortex both from intracranial recordings of neural activity in humans and from recordings of intracortical spiking activity in mice. This was accomplished, both, for an experimental design where the stimulus-predictability was manipulated (in the human brain data) and for one without controlled stimulus-predictability (in the mouse data). We predominantly observe an enhancement in information transfer with increasing neural predictability, suggesting a greater presence of a reliability-coding strategy.

## 2 Methods

### 2.1 Experimental data

We demonstrate the application of the proposed local information dynamics framework on intracranial EEG data and on spike train recordings from the visual system. The first dataset is human brain activity from intracranial electroencephalography (iEEG). The second dataset consists of visually-evoked responses of neurons in mouse visual cortex from the Allen repository brain-observatory [11].

#### 2.1.1 LFP - Intracranial EEG

##### Electrophysiology

We implemented an information-theoretic analysis on intracranial EEG recording of 9 epileptic patients (4 male, 1 left-handed, mean age 31.33 years, range 19-65 years) who were pharmacologically resistant to previous therapies and underwent invasive monitoring using intracranial EEG. The protocol followed community guidelines [12] and was approved by the ethics committee of the University Medical Center Göttingen (protocol number 15/10/18). Patients gave written informed consent to participate in the experiment and were informed that the involvement in the experiment will not interfere in their clinical care. They were monitored constantly by clinical staff and could pause or withdraw from experiments at any time. Patients received financial compensation for their participation. For each patient, brain activity was recorded at bedside from 70 to 128 channels using depth electrodes (DIXI Medical MICRODEEP^®^) with an inter-contact distance of 3.5mm that were stereotactically implanted. The implantation sites and electrode/contact numbers were solely decided based on the clinical needs of each patient. The acquisition system was a Natus telemetry system with a Quantum amplifier.

##### Task

Patients were engaged in a statistical learning task following [13, 14], in which they learned associations between names and faces. The experiment consisted of 2 phases: a learning phase in which a structured stream of names and faces was presented to establish name-face associations, and a test phase were name-face associations were probed. Only data from the test phase are reported here.

In the learning phase, patients were presented with a sequence of names and faces that was ordered into name-face pairs (8 blocks of 108 stimuli pairs). Faces were generated in FaceGen (v3.5.3, Singular Inversions), converted to black and white and luminance histogram equalized using SHINE [15]. Nine unique faces were each paired with a different three-letter name (e.g., “Eva”) that served as a predictor for the face. Names were either spoken or written. Stimuli were presented for 250 ms against a grey screen with a 250 ms inter-stimulus interval. To ensure sustained attention to the stimuli, patients performed a 1-back cover task, which consisted of detecting rarely occurring stimulus duplicates.

During the testing phase, we presented the trained name-face pairs as well as unpredicted conditions in which a name was either paired with a different image of the same identity, a different identity, or a completely novel identity. Only the former two conditions were considered here and combined for analyses. Subjects were instructed to fixate at a fixation cross at the center of the screen. On each trial, first, the predictor — either a written or spoken name — was presented for 250 ms (Fig 1B). Subsequently, an image of a face was presented for 250 ms. Subjects had to indicate by a button press whether the face belonged to the name according to the previously learned associations. The inter-trial interval was randomized between 500 and 1500 ms. Each subject perform 180 trials.

**Figure 1.**
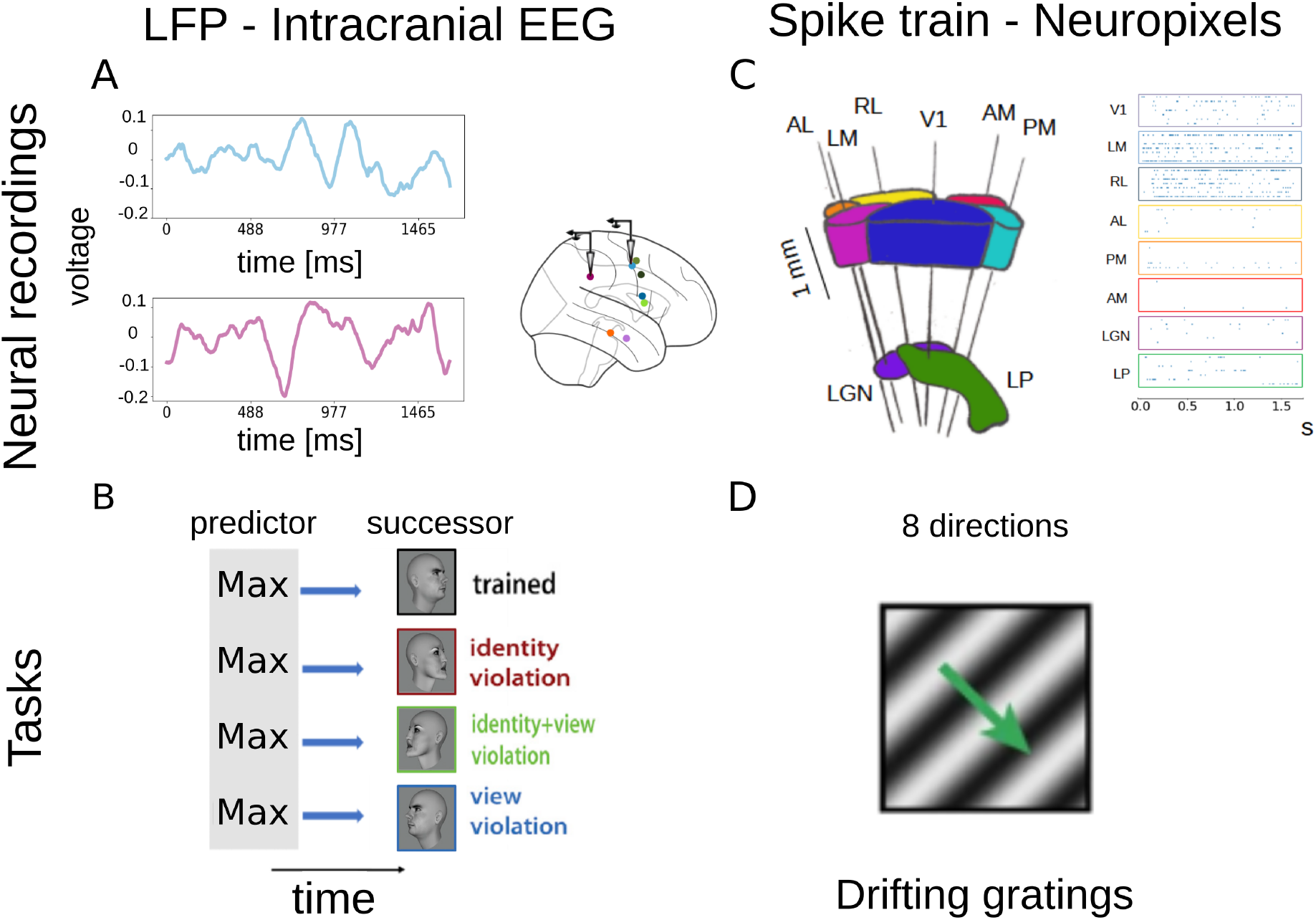
Experimental recordings and tasks for the two datasets. Left panel corresponds to the LFP data, right panel to the spike train. Top panel to the neural recordings, bottom panel to the tasks. **A** We analyzed LFP data from intracranial EEG recording on 9 epileptic patients performing a face processing task. **B** The task involved reading or listening to a given name (e.g., Hardy), which provided the context (i.e., the predictor) for the coming face (i.e., the successor). Subjects were trained on specific name-face pairs to establish the “expected” condition, while subsequent trials introduced changes in either view orientation or face identity to create “unexpected” conditions. **C** We analyzed spiking activity obtained from extracellular electrophysiology recordings of mice brains on visual cortical areas (V1, LM, AL, RL, AM, PM), data obtained from The Allen Brain Observatory database. **D** Recordings in mouse brain were done while presenting a drifting grating visual stimulus. The grating drifts in one of eight directions, at 45° intervals, and at one of five temporal frequencies, ranging from 1 to 15 cycles per second, resulting in 40 distinct grating conditions.

The images were presented on an Asus VG248QE monitor at a comfortable viewing distance. Refresh rate was 120 Hz and a resolution of 1920 x 1080 pixel. Sounds were presented with stereo speakers (Logitech Z200 2.0) mounted on the left and right of the monitor. Responses were collected using a response box (Current Designs) or a keyboard.

##### Data preprocessing

Data preprocessing was performed in Matlab (R2019a, The Mathworks) using the Fieldtrip toolbox (version 20201201, [Fieldtrip Toolbox](https://www.fieldtriptoolbox.org/)) [16]. For information theoretic analyses, data were detrended, re-referenced to a bipolar montage, and decimated to a sample rate of 1024Hz. To identify relevant recording sites, data were bandpass filtered between 0.01 and 30Hz, re-referenced to a bipolar montage, and decimated to 512Hz. Data were epoched from -0.4s to 1.3s relative the onset of the first stimulus (the name).

To identify recording channels with a statistically significant difference between expected and unexpected faces independent of the modality of the preceding name (visual or auditory), we entered the baseline-subtracted, z-scored trial-by-trial data of each recording site into a permutation General Linear Model (GLM, 5000 permutations) as implemented in the Permutation Analysis of Linear Models (PALM) toolbox [17]. Conditions were compared at each time point from the onset of the face stimulus until 500 ms later for the factors modality (auditory/visual) and condition (learned/violation). To determine whether a recording site showed a statistically significant main effect of condition, we used nonparametric combination (NPC) [18] with Tippett’s combining function [19] across the tested time points (two-tailed; irrespective of the sign of the difference) similar to [13]. The resulting p-values were corrected for multiple comparisons across all electrode sites using the False Discovery Rate [20] at q=0.05. Electrodes passing this statistical threshold for the main effect of condition were selected for subsequent analyses.

To localize electrode recording sites, presurgical T1-weighted, anatomical MRIs and postsurgical high density thin-slice computed tomography (CT) were obtained for each patient and co-registered with each other using inhouse scripts. A 3-dimensional reconstruction of each patient’s brain was computed and parcellated using FreeSurfer [21] and brain regions were identified using the Desikan-Killiany atlas [22].

#### 2.1.2 Spiking activity from Allen mouse data - Neuropixels recordings

We implemented an information-theoretic analysis also on spike train recordings from the Allen Brain Observatory database [23]. Data are from the Visual Coding – Neuropixels project [11]. The Visual Coding – Neuropixels project uses high-density extracellular electrophysiology probes to record spikes from the visual cortex. The task consists of a full-field sinusoidal grating that drifts in a direction perpendicular to the orientation of the grating with 8 possible directions (Fig 1D). We select data from ‘brain observatory’, wild type (wt) animals, drifting gratings stimulus, area “area1” and area “area2”. “Area1” and “area2” are selected from the following visual cortical areas: V1, LM, AL, RL, AM, PM. Anatomical hierarchy scores were previously computed to describe projections into and out of these cortical areas via their layer-specific axonal termination patterns (Fig 6A) [24]. Recordings were binned in 3 ms windows.

Using cross-correlation analysis, we selected 40 neuronal pairs from “area 1” to “area 2” that are functionally connected, i.e. those that have the 40 highest values of cross-correlogram (CCG, same as in [25]) average within the time window [-50 ms, 50 ms], and we perform our TE analysis on these pairs. To compute the cross correlation, we perform a shuffling across trials in order to remove correlations that are locked to the stimulus [26], i.e. we compute shuffle-corrected CCGs.

##### Electrophysiology

The Visual Coding – Neuropixels project uses high-density extracellular electrophysiology probes to record spikes from a wide variety of regions in the mouse brain. The experiments were designed to study the activity of the visual cortex. The database used in this work is the “brain observatory” dataset. Data were recorded from 6 possible visual cortical areas: V1, LM, AL, RL, AM, PM (Fig 1C). Examples of the neuronal activity are shown in Fig 1C. We have used the data with the visual drifting grating stimulus (Fig 1D). The cellular responses acquired during the presentation of the drifting grating stimulus provide data to help characterize visual tuning properties of cells, such as orientation tuning (a preference for a grating orientation), direction preference, and temporal frequency tuning (a preference for a specific temporal frequency).

##### Task

From the Allen repository we have used the data during the visual drifting grating stimulus (Fig 1D). In the considered data, the neuronal activity in the mouse visual cortex was recorded while presenting a drifting grating visual stimulus to the animal.

The drifting grating stimulus consists of a full-field sinusoidal grating that drifts in a direction perpendicular to the orientation of the grating. In this experiment, the grating drifts in one of eight directions, at 45° intervals, and at one of five temporal frequencies, ranging from 1 to 15 cycles per second, resulting in 40 distinct grating conditions. The presentation of the visual stimulus is as follows; a grating condition is presented for 2 seconds, and is followed by 1 second of mean luminance gray before the next grating condition is presented. Each of the 40 grating conditions is repeated 15 times, in a random order, and there are intermittent blank sweeps throughout the stimulus (i.e. the 2 second grating condition is replaced with 2 seconds of mean luminance gray). In order to circumvent the non-stationary of the data that is created by the onset response to the visual stimulus, we consider only data from 300 ms to 2 seconds after stimulus onset, thus skipping the onset response; therefore, our data length was 1.7 seconds. In the following we analyzed these data after converting them from spike timestamps (the times when the spikes occurred) to time-binned data using a bin of 3 milliseconds (dt=3 ms).

### 2.2 Mutual information

The mutual information (MI) between two random variables, *X* and *Y*, measures how much information the observation of *Y* provides about *X*, and vice versa (Eq 1). It takes a null value, *I*(*X* : *Y*) = 0, in case the two variables are independent from one another, and *I*(*X* : *Y*) *>* 0 in case of a dependency between *X* and *Y* .

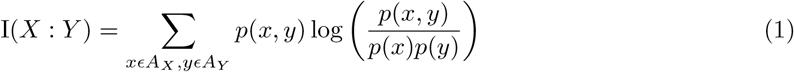

Where *xϵ A*_*X*_ and *y ϵ A*_*Y*_ stand for individual realizations of the random variables *X* and *Y*, and where the fraction inside the log is 1 for all possible joint realizations of two independent random variables.

Mutual information as expressed by Eq 1 measures the information that one random variable has about the other on average. However, the localized counterpart of mutual information (Eq 2) enable us to measure the information provided by single realizations of the random variables *X, Y* .

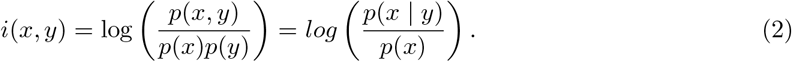

Based on local MI we proceed by describing the local expression of two information-theoretic measures, namely Active Information Storage and Transfer Entropy.

### 2.3 Local active information storage as a proxy of predictability

Using the expression of local mutual information previously defined, we can quantify for a random process — e.g. neural activity — how much residual uncertainty there is at its present moment *t* after observing its past, and also how much uncertainty has already been removed by observing its past; this latter quantity is the mutual information between past state and present sample. We can denote the random process with a sequence of random variables ***X*** = *{X*_*t*_*}* that evolves in time with realizations *x*_*t*_ *∈* ***A***_*X*_*t* . We quantify this uncertainty by estimating the mutual information between the present of the random process, *X*_*t*_, and its own past, **X**_*<t*_.

This measure has been define as Active Information Storage (AIS) of a process ***X*** [8] and quantifies the amount of information stored in the past of ***X*** actively used to predict its next time step. The localized form of AIS (LAIS), i.e. the one computed between single realizations, is defined in Eq. 3.

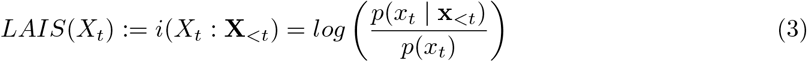

Where *X*_*t*_ stands for the random variable at time *t*, **X**_*<t*_ for the collection of random variables containing the past of *X*_*t*_, and *x*_*t*_, **x**_*<t*_ for single realisations of *X*_*t*_ and **X**_*<t*_ (Fig 2 B).

**Figure 2.**
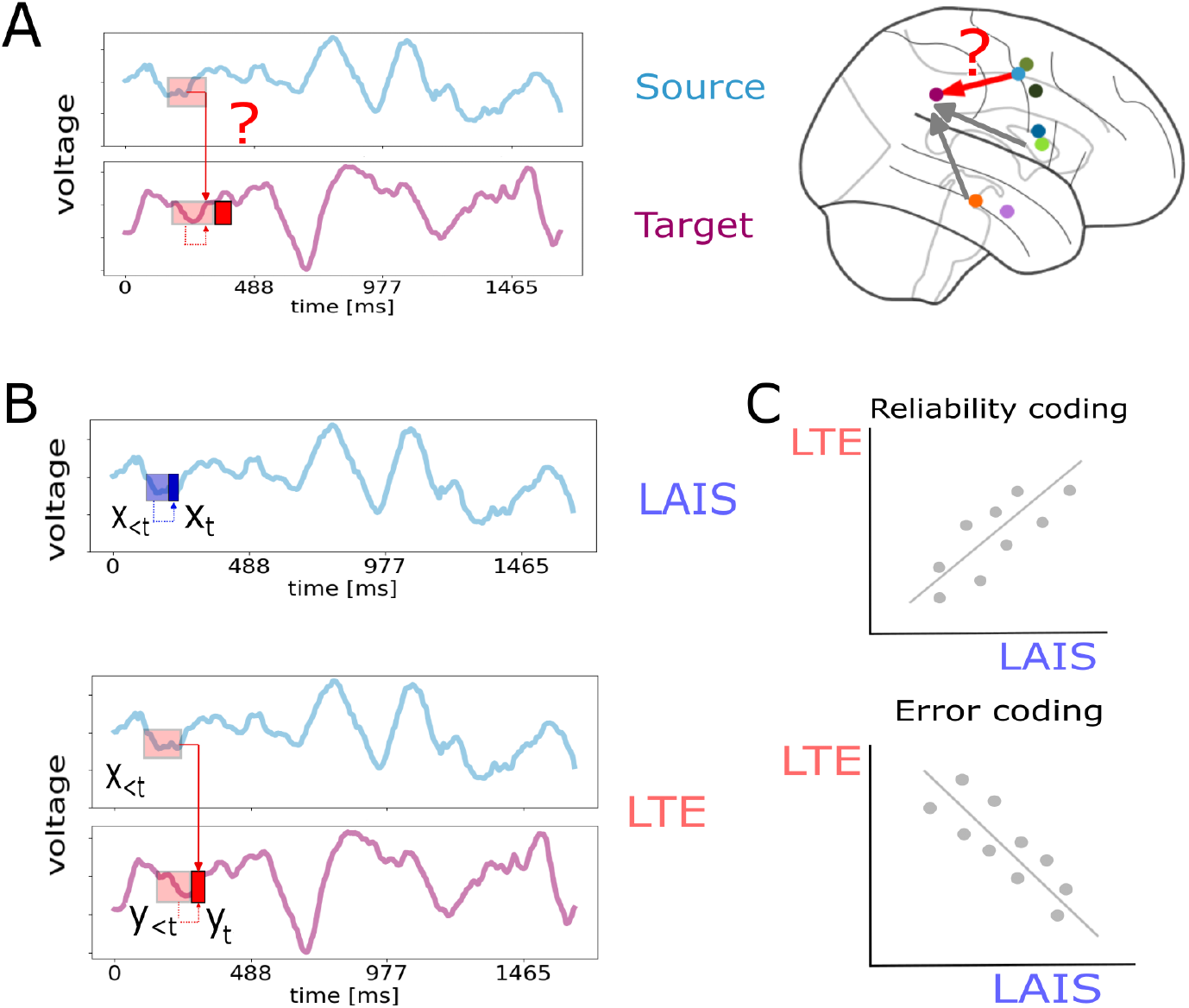
Schematic representation of the steps followed to compute the local storage-transfer correlations (LSTC). **A** We used TE to detect which recorded regions (channels in the iEEG data or units in the extracellular electrophysiological recordings) were connected. **B** For each pair of source-target connections, we computed the local active information storage (LAIS) from the neural activity of the source region and the local transfer entropy (LTE) to the target region. **C** We calculated the LSTC as the correlation between LTE and LAIS, where a positive correlation would signal a reliability coding strategy whereas a negative one an error-coding strategy.

We used LAIS to quantify the predictability of neural activity over time represented by an increase in LAIS. This results in a correspondence between an increase in LAIS – meaning more information is stored in the past and used to predict the next time step – and an increase in the predictability of the neural activity. In order to capture all the relevant past that contributes to the prediction of the next time step, it is necessary to quantify the contribution of semi-infinite window in the process’s past [27]. However, given the computational complexity of taking all the past states, we worked under the approximation of markovian processes with finite memory. This assumes that the probability of a state conditioned on its past within a finite time window, equals its probability conditioned on all its semi-infinite past (Eq 6 in section 2.7.1).

In the context of predictive coding, we interpret a decrease in LAIS representing a decrease in predictability, as a signature of a mismatch between expected and actual neural response, i.e., a prediction error. On the contrary, an increase in the predictability would signal a match between the expected and the actual neural response. In order to study the communication of either predictable or unpredictable states, we describe in the following section an information-theoretic measure of information transfer.

### 2.4 Local transfer entropy as a measure of information transfer

Local transfer entropy (LTE) captures the information transfer from source process(es) — e.g., neuron or brain area — to a target process — e.g., another brain area. We denote the random source process as ***X*** and the random target process as ***Y*** . Information transfer is represented as a reduction in uncertainty at time point *Y*_*t*_ after observing the past of ***X***, *given that we know the past of* ***Y***.

In other words, local transfer entropy [28] quantifies a reduction in uncertainty that was not already possible by observing the past of ***Y*** (Eq 4). Note that this reduction in uncertainty is due to information about the current state of the target uniquely provided by the past of the source, and the information about the current state of the target synergistically provide by both, the past of the source, and the past of the target.

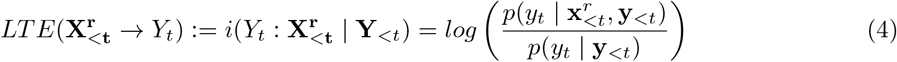

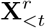 stands for the collection of random variables containing the past of the source processes and the superscript *r* for the number of source processes. *Y*_*t*_ stands for the random variable at time *t*, **Y**_*<t*_ for the collection of random variables containing the past of *Y*_*t*_. *y*_*t*_, **y***<t* and 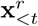 stand for realisations of *Y*_*t*_, **Y**_*<t*_ and 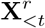, respectively.

Before we estimate the local transfer entropy at each time step, we have to identify from a given set of candidate sources which are the actual sources that transfer information to the target. In this work, we used the average transfer entropy (TE) (Eq. 5) as a measure of brain connectivity.

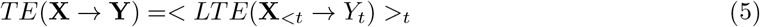

Just for clarity, in this expression we dropped the superscript *r* given that we are only looking at the transfer entropy between one source process ***X*** to a target ***Y*** . Here, we used both a bivariate and multivariate approach of transfer entropy to identified neurons or brain areas connected.

### 2.5 Connectivity analysis

#### 2.5.1 Multivariate transfer entropy analysis of iEEG recording

We used a multivariate measure of transfer entropy to infer brain connectivity (Fig 2A) from intracranial EEG recordings of human epileptic patients (Fig 1A) [29]. The multivariate approach allowed us to study the interaction between two given brain areas in the context of the rest of the analyzed set. The implementation of a multivariate analysis was feasible due to the small number of analyzed channels per subject — between 6 to 9 channels. We analyzed only those channels that presented a statistically significant difference between expected and unexpected stimuli — see 2.1.1 for a detailed description of the task and the procedure for the channel’s selection. However, for the transfer entropy connectivity analysis we used both conditions, i.e., expected (predictable) and unexpected (unpredictable). The use of TE as a connectivity measure provides a directionality for the pair of connected brain areas. Thus, for each pair of connected areas we will obtain a source and target brain areas.

#### 2.5.2 Bivariate transfer entropy analysis of spike train data

For spike train recordings of the mouse visual cortex, we inferred the connectivity between neuronal pairs across distinct brain areas using a bivariate transfer entropy approach. In contrast to the iEEG recordings, which included only a limited number of recording channels, the spike train data contained an average of 80 neurons recorded per brain area, and multiple brain areas, therefore thousands of neuronal pairs were available. This large number of neuronal pairs makes a multivariate approach impractical for the spike train data, as the multivariate transfer entropy estimation algorithm (described in section 2.7) scales cubically with the number of processes analyzed in the worst case (see Supporting Information of [30]). However, in the bivariate case, this limitation is addressed by analyzing neuronal pairs individually, corresponding to the case with r=1 in Eq 4. To optimize the bivariate analysis, we selected neuronal pairs based on a preliminary cross-correlation analysis (see Section 2.1.2).

### 2.6 Local Storage Transfer Correlations

We studied the information dynamics unfolded for each connection (between brain areas in the LFP data and between neuronal pairs in the spike train data) that showed a significant transfer entropy. The goal of this analysis is to detect the presence of predictive processing and the type of coding strategies — i.e., coding for reliable or error information. We studied the presence of predictive processing at the cortical level by the detection of *local storage-transfer correlations* [10]. For a given connection, the *local storage-transfer correlations* (LSTC) are defined as the correlation between the LAIS — predictability — of neural activity of the source brain area or neuron with the LTE — information transfer — to the target brain area or neuron.

Through the sign of these correlations between predictability and information transfer we are able to tease apart two types of coding strategies related with predictive processing theories. A positive sign for the LSTC would indicate an increase in the information transfer to the target of the connection when the predictability of the source increases. Thus, a positive correlation would be a signature of coding for reliable information (Fig 2C). On the contrary, a negative correlation would indicate a suppression in the information transfer when the predictability of the source increases. Then, a negative correlation would be a signature of error coding (Fig 2C).

The LSTC analysis was implemented on every cortico-cortical connection detected on both iEEG recordings of epileptic patients and spiking activity of mouse visual system. Importantly, due to the adoption of a neuro-centric perspective that relies on quantifying predictability directly from neural activity, the LSTC analysis of the iEEG recordings incorporated both experimental conditions (i.e., expected and unexpected).

### 2.7 Estimation of information-theoretic measures

In this study we will estimate LAIS and LTE from intracranial EEG recordings (2.1.1) and from extracellular electrophysiological recordings (Allen data [11]). For the intracranial EEG recordings we will use the multivariate Transfer Entropy algorithm, for the extracellular electrophysiological data (spike trains) we will analyze the data using the bivariate Transfer Entropy because of the scarcity of relevant links between neurons.

#### 2.7.1 Intracranial EEG recordings

Both the definition of LAIS (section 2.3) and LTE (section 2.4) require the estimation of the probability distributions that underlies the observed data.

For continuous random variables one can estimate them using the KSG estimator [31]. In sections 2.3 and 2.4 we showed that both information-theoretic measures are based on (conditional) mutual information (MI). KSG is an estimator of MI from k-nearest neighbor statistics. It consists of taking the distance between a realization of the random variables and its k-nearest neighbor in the high-dimensional joint space of random variables. This distance is projected to the low-dimensional marginal spaces of the random variables and the number of realizations that fall within this distance are counted. We used k=8 as the number of neighbors to search in the high-dimensional joint space. We applied the KSG estimator on the LFP data obtained from the intracranial EEG recordings.

To estimate LAIS and LTE from continuous random variables it is necessary to estimate the MI between the present and past states of the random processes. This past embedding has to take into account the appropriate lags to map the time series of the random processes into trajectories in a state space.

In theory, to build this embedding it should take into account all the past of the random process but we worked under the assumption of Markovian processes with finite memory *l*_*M*_ . The Markovian assumption implies that the present of the system *X*_*t*_ is independent of the past variables before the maximum lag *l*_*M*_ given that we observed the past until *l*_*M*_ (Eq. 6).

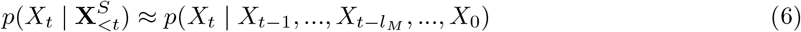

Where 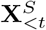 contains selected random variables in the past of the random process X until *l*_*M*_ .

We used a nonuniform embedding approach [32] to select the time lags that were most informative — i.e, with the highest mutual information — from the set candidate time lags 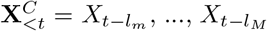. The nonuniform embedding consists of selecting those random variables in the past of *X*_*t*_ from a set of past candidate variables 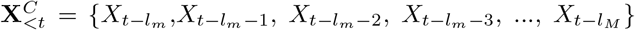 whose mutual information with *X*_*t*_ is the highest. The set 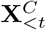 differs from 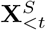 in that 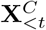 contains all the random variables in the time window [*t−l*_*m*_, *t−l*_*M*_] whereas 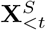 just the ones that are maximally informative. We handle the computational complexity given by the size 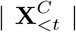 of the candidates set by making use of a greedy forward-selection algorithm. This algorithm consists of maximizing the information contained in the variable set 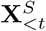 with respect to *X*_*t*_ by using conditional mutual information (CMI) as selection criterion,

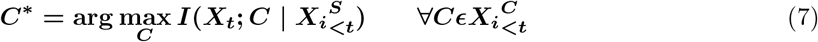

where 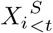 is the set of variables already selected in the *i* th step of the algorithm and C are candidate variables from the set of candidates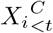. Note that in the candidates set there is also the subscript *I* indicating that the candidates set also depends on the iteration of the algorithm. After selecting the *C*^*∗*^ a surrogate test is used to assess that 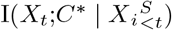 is statistically significant. If *C*^*∗*^ is significant then both sets 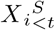 and 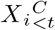 are updated by removing the *C*^*∗*^ from the candidates set 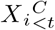 and adding it to the set of selected sources 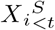 .

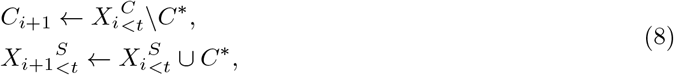

After building the set 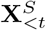we computed LAIS as defined in Eq. 3 with **X**_*<t*_ only containing the most informative random variables in the past.

To estimate LTE, we build the set of target’s past 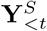 of most informative lags, and afterwards we select the most informative lags from the set 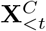 of past candidate variables of the source processes.

We proceed iteratively as before by selecting *C*^*∗*^,

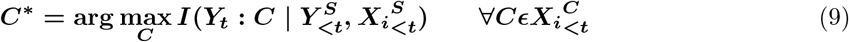

In this way, we incorporate only those random variables in the source’s past that are the most informative *conditioned* on the target’s past.

Both the estimation of bivariate and multivariate TE was done with the same iteration procedure. In Eq. 4 we mentioned that 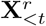 stands for the contribution of all sources. Similarly, here 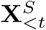 contains all the selected random variables in the source’s past. The multivariate analysis implemented for the intracranial EEG data implied that the set of candidate variable of the source’s past 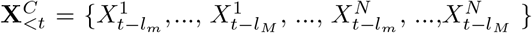 contains all the random variables within the window [*t − l*_*m*_, *t − l*_*M*_] for the *N* source processes.

In the analysis of intracranial EEG data, we estimated LAIS using a time window between *l*_*m*_=1 ms to *l*_*M*_ =15 ms in the past. For the estimation of LTE, we looked in the target’s past also within a time window between *l*_*m*_=1 ms to *l*_*M*_ =15 ms, and in the source’s past between *l*_*m*_=3 ms to *l*_*M*_ =30 ms.

We used a software implementation of the described approach provided by the IDTxl Python toolbox [33], which internally makes use of the KSG estimators implemented in the JIDT toolbox [34]. A more detailed description of the algorithm here explained can be found [30], which includes an explanation about a hierarchical statistical testing structure to handle the family-wise error rate of the repeated testing during the iteration over the candidates set.

#### 2.7.2 Spike train data

We applied the analysis of local active information storage and local transfer entropy, and their correlation also to data from spike train recordings from the Allen Brain Observatory database [11]. As described in the previous section, the estimation of local information-theoretic quantities requires the estimation of probability distributions that underline the observed data. In this section we describe the estimation of conditional mutual information (MI) from spike data. We follow the procedures already used in [10]. The most straightforward approach to estimating MI from discrete data (Eqs 3 and 4) is by replacing probability mass functions by the relative frequencies of symbols observed in the data [35]. These so-called “plug-in estimators” are well-known to exhibit negative bias for finite data, for which analytic bias-correction procedures exist [36, 37]. These bias-correction approaches, formulated for non-local variants of mutual information, may be adapted for the use with localized measures to obtain locally bias-corrected estimators of LAIS and LTE (see S1 Text in [10]). Furthermore, statistical testing against estimates from surrogate data may be applied to handle estimator bias by treating the estimate of information transfer or storage as a test statistic to be compared against a Null-distribution generated from estimates based on suitable surrogate data. Before applying conditional mutual information estimators, the relevant past states of the time series involved have to be defined. As explained in the previous section, both AIS and TE quantify the information contained in the semi-infinite past of a time series up to time point t; however, most information is most often contained within the immediate past of the present sample, *X*_*t*_ [38, 39]. Hence, we define an “embedding” of the past of the time series, 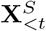, i.e., a collection of past variables up to a maximum lag, selected such the embedding is maximally informative about *X*_*t*_.

For our analyses the recorded spike trains were binned into 3 ms segments. We optimized nonuniform past-state embeddings for each cell pair recording using the greedy algorithm implemented in [33]. For spiking data we estimated LAIS and LTE by searching for the most informative random variables in a time window from the previous *dt* to a certain maximum time lag in the past. Specifically, we estimated LAIS using a time window between *l*_*m*_=1 ms to *l*_*M*_ =50 ms in the past (with data binned in 3 ms windows). For the estimation of LTE, we looked in the target’s past also within a time window between *l*_*m*_=1 ms to *l*_*M*_ =50 ms, and in the source’s past between *l*_*m*_=3 ms to *l*_*M*_ =60 ms.

These lags assume that source spikes with an inter-spike interval (ISI) of 10-20 ms are relevant for triggering a spike from one visual cortical area to another one (Figure 3C and Extended Data Figure 7 in [40]). Our estimate of the necessary maximum delays and time lags was based on evaluating the lagged cross correlations within pairs and selecting lags and delays such that the peak of the lagged correlations would be covered. This selection was necessary to keep the computational burden manageable which scales quadratically with the number of investigated lags.

**Figure 3.**
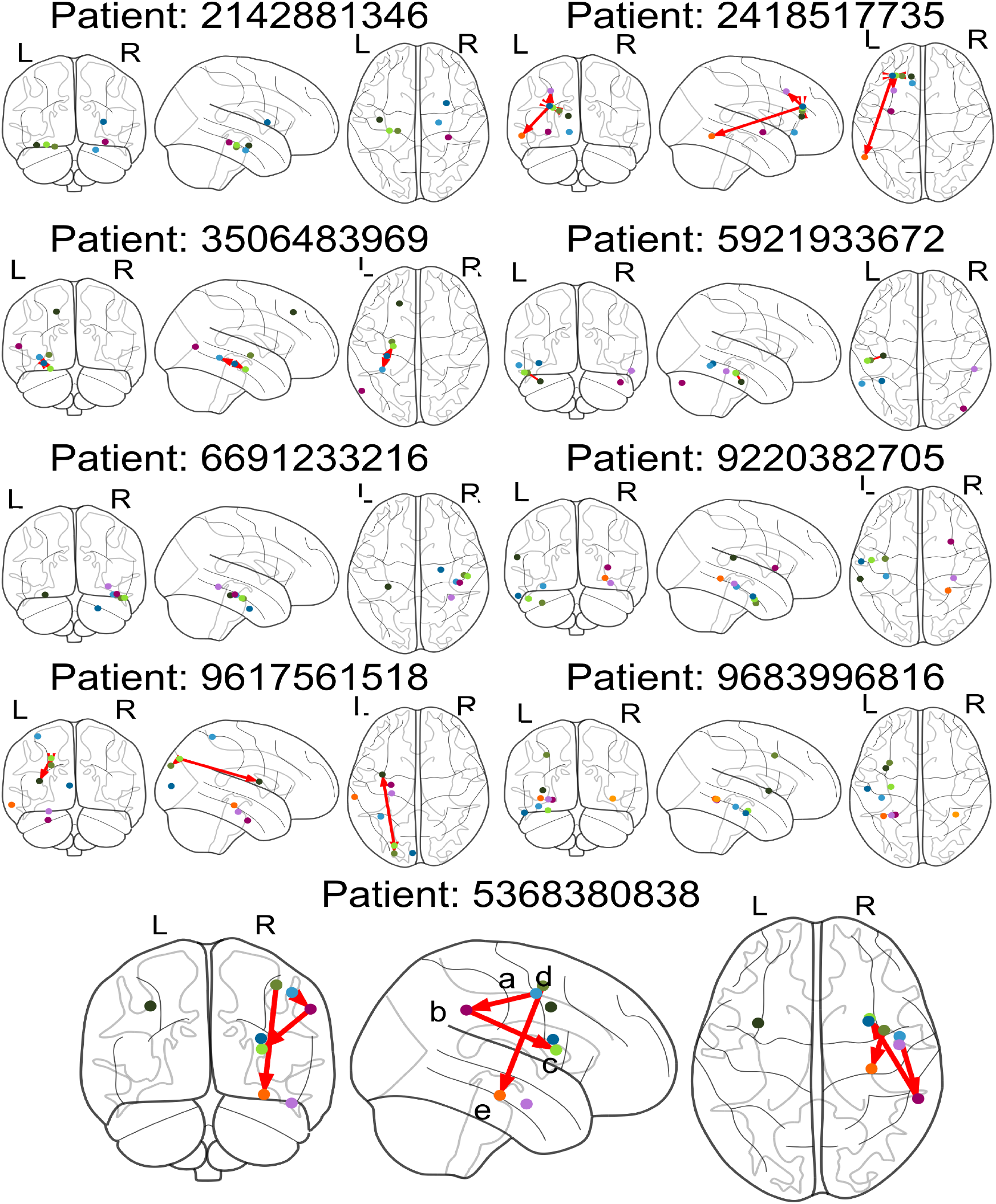
Cortico-cortical multivariate transfer entropy network for each patient. The colored dots represent the locations of iEEG recording sites included in the TE network analysis. The red arrows stand for the transfer entropy links. We observed at least one significant link (p *<* 0.004) in 6 out of 9 patients — patients 2142881346, 9220382705 and 9683996816 did not present any significant link and were therefore excluded from the Local Storage-Transfer Correlations analysis. Significant connections are found in frontal, parietal and temporal areas. As an exemplary subject (bottom) we observed for patient 5368380838 cortical information transfer from right rostral middle frontal (a) to right supramarginal (b), from right supramarginal (b) to right insula (c) and from right caudal middle frontal (d) to right hippocampus (e).

After building the set 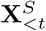 we computed LAIS as defined in Eq. 3 with **X**_*<t*_ only containing the most informative random variables in the past.

To estimate LTE, we build the set of target’s past 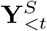 of most informative lags, and afterwards we select the most informative lags from the set 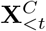 of past candidate variables of the source processes. We proceed iteratively as before by selecting *C*^*∗*^, using the same formula as in Eq. 9.

For the calculation of LTE and LAIS we used the software already cited: the IDTxl Python toolbox [33], which for the spiking data internally makes use of the discrete estimator implemented in the JIDT toolbox [34].

Subsequently, we computed the bias following the procedure in S1 Text [10](Supporting information *Localized bias Treves-Panzeri correction*) to obtain the bias-corrected estimators of LTE and LAIS.

## 3 Results

### 3.1 Transfer Entropy analysis iEEG recordings

We performed a network analysis of intracranial EEG recordings to assess the information transfer between cortical areas. To this end we used a multivariate implementation of transfer entropy as a measure of connectivity. This measure allowed the detection of links where a source brain area transfers information to target area, in the context of the rest of the analyzed cortex. This analysis revealed 17 cortico-cortical connections with significant transfer entropy (p *<* 0.004, see [30] for a detailed explanation of the hierarchical statistical test implemented) across all patients (Fig 3) encompassing frontal, parietal and temporal areas. The analysis was implemented for each patient individually, showing in 6 out of 9 patients at least one significant link.

For each cortico-cortical connection we studied the information dynamics by means of local transfer entropy and local active information storage. These measures were used to study whether the information transfer to the target area increases or decreases with the predictability of the source area.

### 3.2 Local Storage-Transfer Correlations on iEEG

We studied the information dynamics unfolded for each detected cortico-cortical connection to assess the presence of predictive processing and the type of coding strategy — i.e., reliability or error coding strategy. For this purpose, we computed for each link the Local Storage Transfer Correlation (LSTC) (2.6), that is the correlation between LAIS and LTE. We quantified both LAIS and LTE, incorporating both types of recorded conditions (i.e., expected and unexpected), given that our neuro-centric approach relies on obtaining neural predictability directly from neural recordings (see S1 Text for a study on the sensitivity of TE to stimulus predictability).

We found that all 17 links presented significant correlations (p *<* 0.05, Bonferroni corrected for 17 links) between LAIS and LTE (Table 1) with a positive sign ranging between r=0.006 and r=0.093 (Fig 4B). For an exemplary patient (Fig 3, patient 5368380838) we observed from right caudal middle frontal to right hippocampus a correlation of r=0.055, from right rostral middle frontal to right supramarginal r= 0.093 and from right supramarginal to right insula r= 0.024 (Fig 4A).

**Table 1.**
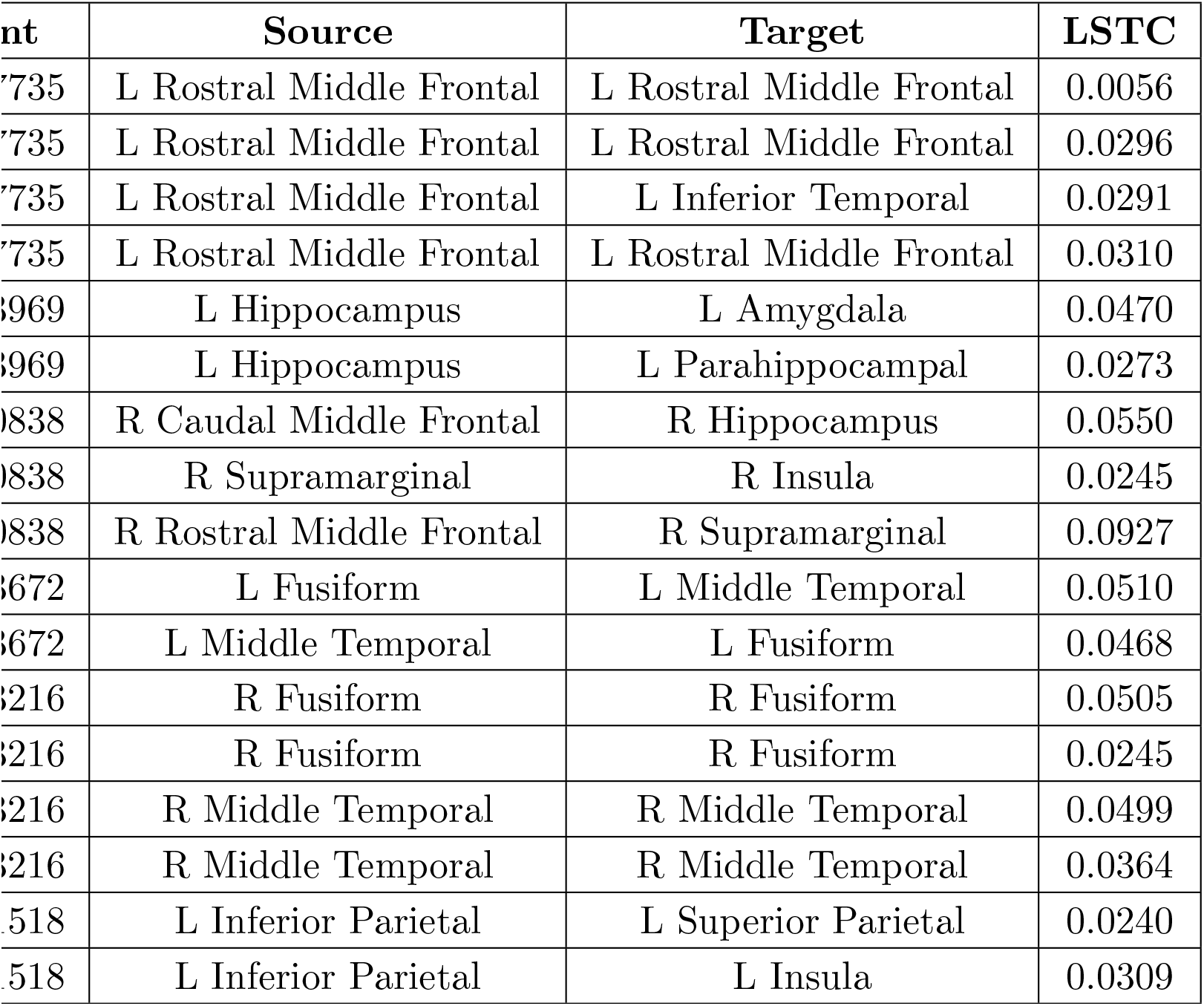
Cortico-cortical connections with significant transfer entropy (p *<* 0.004) for the human intracranial EEG recordings. The LSTC column shows the significant local storage-transfer correlations (p*<*0.05, Bonferroni corrected). The whole set of brain regions included in the TE analysis can be observed in Fig 3. R, right; L, left.

| Patient | Source | Target | LSTC |
| --- | --- | --- | --- |
| 2418517735 | L Rostral Middle Frontal | L Rostral Middle Frontal | 0.0056 |
| 2418517735 | L Rostral Middle Frontal | L Rostral Middle Frontal | 0.0296 |
| 2418517735 | L Rostral Middle Frontal | L Inferior Temporal | 0.0291 |
| 2418517735 | L Rostral Middle Frontal | L Rostral Middle Frontal | 0.0310 |
| 3506483969 | L Hippocampus | L Amygdala | 0.0470 |
| 3506483969 | L Hippocampus | L Parahippocampal | 0.0273 |
| 5368380838 | R Caudal Middle Frontal | R Hippocampus | 0.0550 |
| 5368380838 | R Supramarginal | R Insula | 0.0245 |
| 5368380838 | R Rostral Middle Frontal | R Supramarginal | 0.0927 |
| 5921933672 | L Fusiform | L Middle Temporal | 0.0510 |
| 5921933672 | L Middle Temporal | L Fusiform | 0.0468 |
| 6691233216 | R Fusiform | R Fusiform | 0.0505 |
| 6691233216 | R Fusiform | R Fusiform | 0.0245 |
| 6691233216 | R Middle Temporal | R Middle Temporal | 0.0499 |
| 6691233216 | R Middle Temporal | R Middle Temporal | 0.0364 |
| 9617561518 | L Inferior Parietal | L Superior Parietal | 0.0240 |
| 9617561518 | L Inferior Parietal | L Insula | 0.0309 |

**Figure 4.**
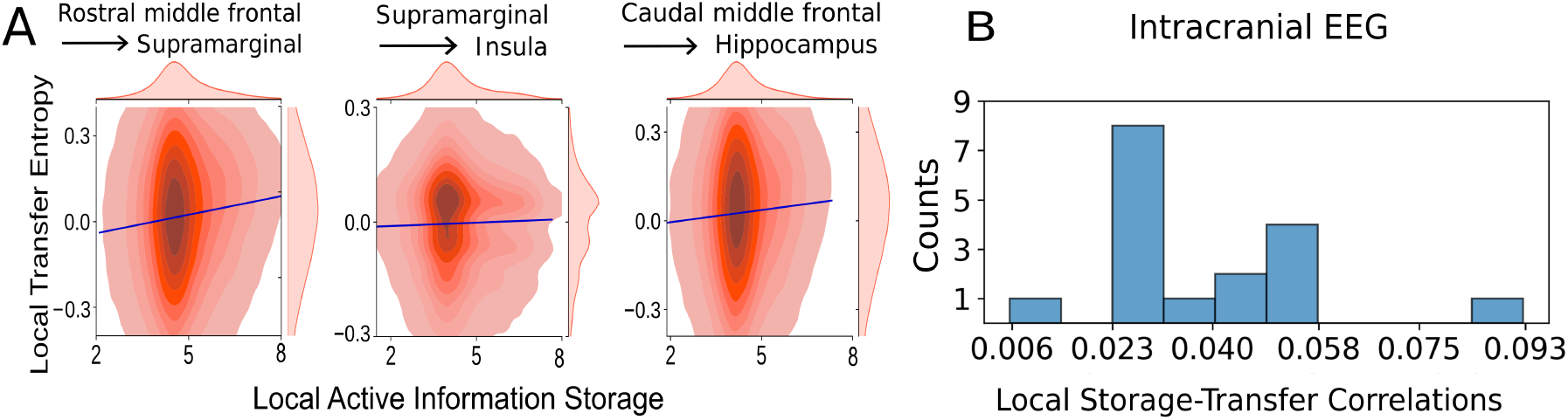
Local storage-transfer correlations for the human intracranial EEG recordings. **A** LSTC for the three cortico-cortical connections of the exemplary patient 5368380838 (bottom Fig 3). **B** Histogram of LSTC for all 17 cortico-cortical connections across the entire cohort.

Across all subjects, our results showed that frontal, temporal and parietal regions are involved in the transfer of information (Table 1). However, the observed connections did not account for the same regions for all patients. This is due to the high sparsity of intracranial EEG recordings on epileptic patients, since the electrodes’ location must be clinically driven [12].

### 3.3 Local Storage-Transfer Correlations on spike data

To assess whether LSTCs are potentially detectable at the level of multiple single-unit recordings, we also analyzed data from mouse cortex provided by the Allen Brain Institute. These data pose several additional challenges compared to the analyses of LSTCs in iEEG. First, information transfer between individual cells will be hard to detect, as monosynaptically connected cell pairs may not be present in the dataset, and we therefore may analyze information relayed across multiple cells; second, cortical neurons typically receive a large number (100-1000) of different inputs that are processed together to form the receiving neuron’s output. The influence of a single input neuron on this output will be small and hard to detect. Given these challenges we consider the results presented below only as a first exploratory study of the potential to analyze LSTCs in cortical single-unit recordings.

#### 3.3.1 Transfer Entropy analysis on spike data

We considered all possible directed connections between a source brain area and a target brain area among these 6 cortical brain areas: V1, RM, LM, AL, PM, AM. For each given source and target area, we selected the first 40 neuronal pairs (source neuron and target neuron) that display the highest pairwise lagged correlation. For each brain area pair, we found a number *m* of neuronal pairs whose TE are significant, i.e. we found a significant TE only on some of the 40 pairs analyzed. For the *m* values for each brain area pair see Table 2. On average we found 9 pairs between two brain areas that showed a significant TE. On each of these pairs we further computed the LAIS and LTE (described in Section 2.7.2).

**Table 2.** LSTCs with relative p-value for all the pairs analyzed (pairs with a significant TE), sorted by source brain area and target brain area. The information is transferred from the source brain area to the target brain area. Each entry represents the significant LSTC value found for one neuronal pair out of the *m* pairs that display a significant TE. NS: LSTC values that were not significant. NA: no pairs with a significant TE was found.

| Source Area<br>Target Area | V1 | RL | LM | AL | PM | AM |
| --- | --- | --- | --- | --- | --- | --- |
| **V1** |  | NS<br>(m=14) | NS<br>(m=15) | -0.0101<br>p=0.002<br>(pair 3)<br>(m=8) | NS<br>(m=3) | +0.0098<br>p=0.001<br>(pair 25)<br>(m=14) |
| **RL** | NS<br>(m=19) |  | NS<br>(m=9) | NS<br>(m=11) | +0.010<br>p=0.001<br>(pair 31)<br>(m=7) | NS<br>(m=16) |
| **LM** | NS<br>(m=6) | NS<br>(m=3) |  | NS<br>(m=10) | NS<br>(m=4) | NS<br>(m=18) |
| **AL** | -0.0086<br>p=0.007<br>(pair 28)<br>(m=7) | NS<br>(m=8) | NS<br>(m=7) |  | NS<br>(m=2) | NS<br>(m=7) |
| **PM** | +0.0082<br>p=0.007<br>(pair 32)<br>+0.0097<br>p=0.003<br>(pair34)<br>(m=4) | NS<br>(m=9) | NS<br>(m=8) | +0.0092<br>p=0.005<br>(pair 7)<br>(m=3) |  | +0.0081<br>p=0.001<br>(pair 25)<br>(m=8) |
| **AM** | +0.0087<br>p=0.007<br>(pair 12)<br>(m=4) | NS<br>(m=6) | -0.0090<br>p=0.003<br>(pair 29)<br>(m=9) | NS<br>(m=16) | NA<br>(m=0) |  |

#### 3.3.2 Local Storage-Transfer correlations on spike data

We computed the local AIS and local TE with the procedure explained in Section 2.7.2 on the neuronal pairs that display a significant TE between the source neuron and the target neuron. We computed the LSTCs (see definition in 2.6) for each neuronal pair that had a significant TE (see Fig 5 for examples). Fig 5B shows the histogram of the significant values for LSTCs found across all brain areas (p *<* 0.05, Bonferroni corrected per area pair). We report the significant LSTC values per neuronal pair in Table 2. Our results are summarized in the circuit depiction in Fig 6B. We used the anatomical hierarchy as previously derived via the layer-specific axonal termination patterns of projections into and out of these cortical areas [24, 40]. Nine neuronal pairs - from and to different visual areas - showed a significant LSTC. We found 4 significant links to higher areas (PM, AM) that show a positive LSTC, one link from LM to AM that exhibited a negative LSTC. Lower and middle brain areas display both negative and positive LSTCs.

**Figure 5.**
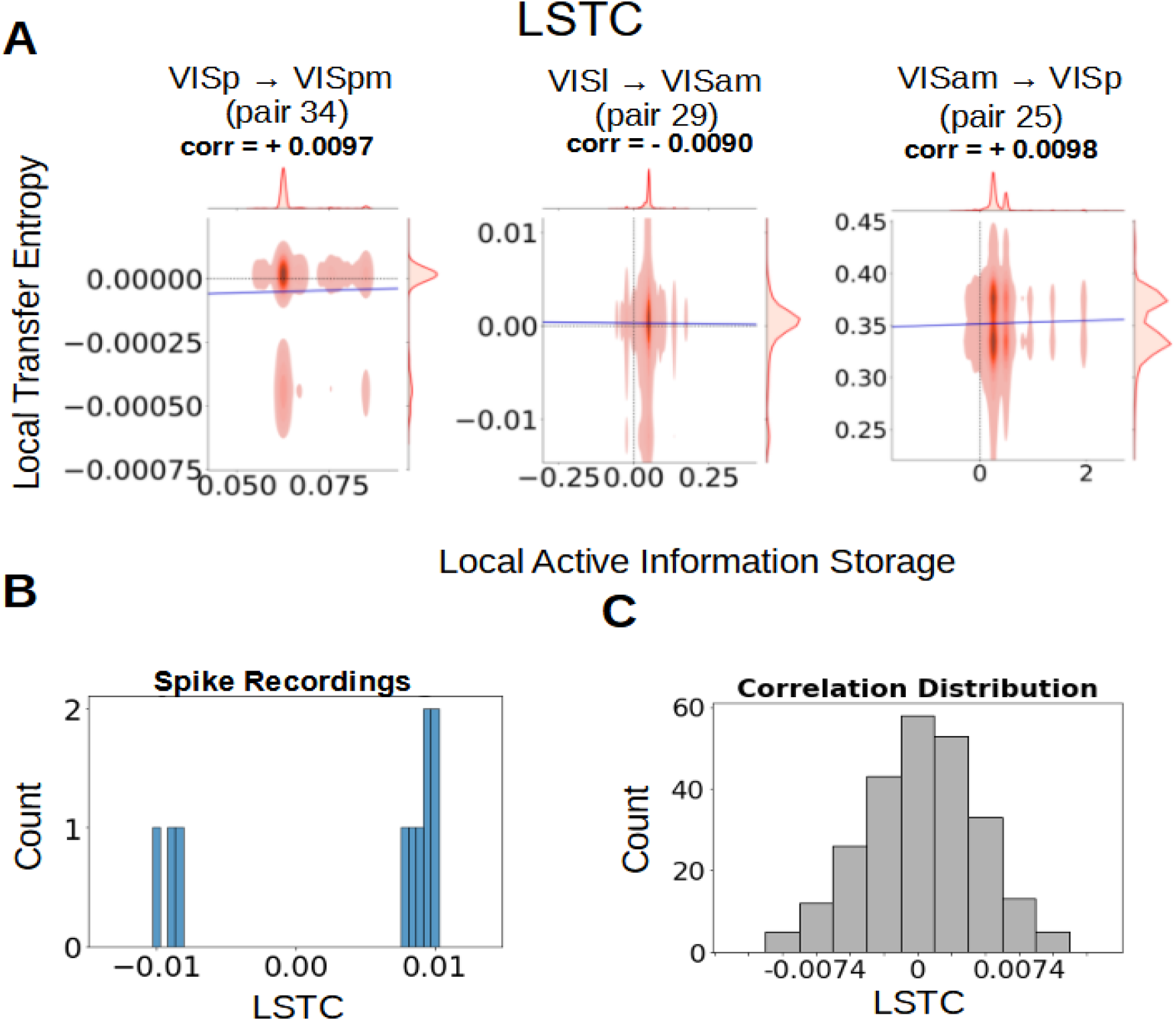
Local storage-transfer correlations for the Allen neuronal recordings. **A** LSTC for some exemplary neuronal pairs that show a significant LSTC. The local transfer entropy is plotted as a function of the local storage information, showing the correlation between the two quantities. For example, the first graph from the left shows the LSTC for the neuronal pair n. 34 from a source neuron in VISp to a target neuron in VISpm. **B** Histogram of LSTC for all neuronal pairs from all source areas to all target areas (possible areas are VISp, VISrl, VISl, VISal, VISpm, VISam) that showed a significant correlation. **C** Histogram of LSTC for all neuronal pairs from all source areas to all target areas (possible areas are VISp, VISrl, VISl, VISal, VISpm, VISam) both non-significant and significant LSTC are included.

**Figure 6.**
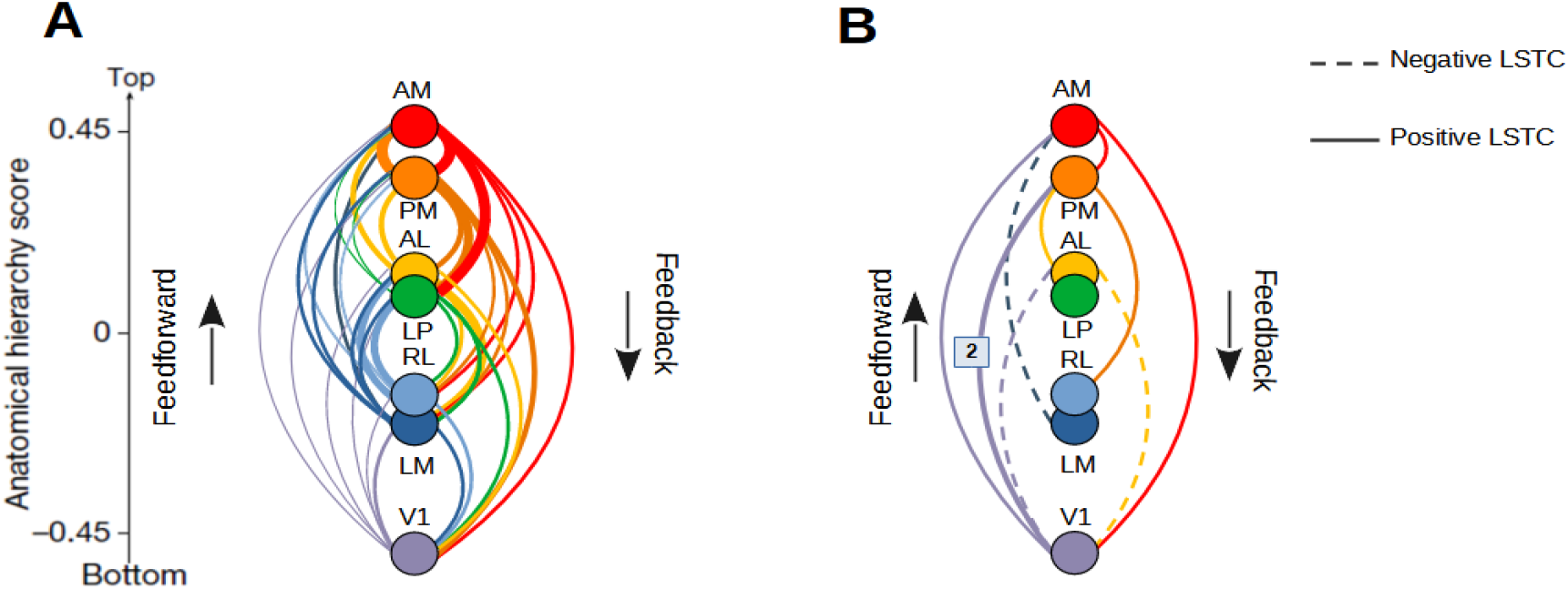
**A** All connections between brain areas that are considered in our analysis. Hierarchy is shown bottom up. Anatomical hierarchy score were previously computed to describe projections into and out of cortical areas via their layer-specific axonal termination patterns [24] and reported in [40]. **B** Area connections in which a neuronal pair with a significant LTSC is found. A full line or a dash line represent a positive correlation value found or a negative correlation value respectively. If not stated otherwise the connection found is relative to a single neuronal pair with a significant TE and a significant LTSC. Two significant pairs are found from area V1 to area PM. Neuronal recordings used are from the Allen repository [11].

### 4 Discussion

We tested the presence of predictive processing (PP) coding strategies at the cortical level using a purely information theoretic framework in order to minimize assumptions imposed by the experimenter. Predictive processing coding strategies differ with respect to the type of information that is passed on between neurons or brain areas, i.e. predictable or unpredictable information; these strategies can can be distinguished information-theoretically by the sign of the correlation of the active information storage and the transfer entropy. The strategy of encoding for reliable information would be present as an enhancement in the information transfer when the predictability of the neural activity increases, i.e., a positive correlation. On the contrary, the error coding strategy would be present as suppressed information transfer when the predictability increases, i.e., a negative correlation.

We found support for the presence of PP in the form of significant correlations between the information transfer and predictability in cortical areas of epileptic patients and provide some tentative evidence for the existence of such correlations in the visual cortex of mice.

More specifically, in coarse-grained activity, i.e. the LFP signal obtained from iEEG recordings, we found an enhancement in the information transfer to the target brain area when the predictability of the source cortical area increased — for cortical connections with significant information transfer.

In contrast, at a finer resolution, i.e. the spiking activity of neurons in the mouse visual system, we did also find decreases in information transfer when the predictability of the source increased, although the opposite correlation was still a more frequent finding.

These findings support that at the cortical level both types of coding strategies are employed as a processing mechanism. However, the evidence for only the reliability coding strategy in the LFP signal, suggests that our predictive processing measure might present a sensitivity to the resolution of neural recordings.

#### 4.1 Cortico-cortical communication seems to predominantly code for reliable information

Cortico-cortical communication presents both types of coding mechanisms, as evidenced by the presence of positive and negative local storage-transfer correlations (LSTC). This measure reflects the correlations between the predictability of neural activity and information transfer across the cortex. We identified significant LSTCs in both LFP data and spiking activity. Specifically, in human iEEG recordings, we detected 17 LSTCs in the LFP signal, while in the spiking activity of neurons in the mouse visual system, we identified a total of 9 significant LSTCs.

Each correlation detected from the iEEG signal exhibited a positive sign, indicating a coding strategy for reliable information communication across cortex. This strategy implies the encoding of predictable information, characterized by an increase in information transfer with predictability.

The positive sign of the LSTC was consistently observed across all cortical regions, including frontal, parietal, and temporal areas. However, we interpret this positive LSTC not as representative of a coding mechanism between two specific brain regions, but rather as evidence of a predominant coding for reliable information over the entire analyzed cortex. This interpretation is suggested by the fact that in our subjects with epilepsy, as necessarily had to record from many different brain regions.

In contrast, spiking activity exhibited putative signatures of both coding strategies. We primarily observed a signature of coding for reliable information, reflected by 66.67% cortical connections with positive LSTC. However, we also observed evidence of an error coding strategy, showed by 33.33% connections with negative LSTC. The latter strategy involves the encoding of unpredictable information, characterized by a decrease in information transfer with predictability.

The observation of positive and negative LSTCs at the level of spike recordings possibly indicates that at this more localized level a coding for unpredictable (error) information may possibly be present. In this case, we hypothesize that the smaller number of connections exhibiting an error coding strategy may explain the absence of negative LSTC in the LFP signal: It is possible that results in coarse-sampled activity, where we only observed positive correlations, may originate from the LFP signal representing an average activity over a population of neurons, where positive LSTCs then simply dominate the average.

However, this study did not systematically explore the sensitivity of the LSTC measure with respect to resolution level in one species and setting. We, therefore, suggest that future research evaluates the LSTC measure in such a context.

This study revealed the presence of LSTC across different resolution levels and under two distinct experimental conditions, adopting a neuro-centric perspective. By quantifying neural predictability, we identified the specific predictive coding mechanism involved in a face-processing task with controlled stimulus-predictability (in human subjects) and in a drifting grating visual task without controlled stimulus predictability (in mice).

#### 4.2 Explaining away at the cortical level

Empirical evidence has revealed two types of biophysical coding strategies, demonstrating both top-down enhancement and inhibition of neural activity. This modulation of the neuron’s activity results in either an increase or decrease in information transfer by the modulated neuron. Specifically, it has been shown that sensory neurons exhibit top-down enhancement of their activity when encoding task-relevant stimulus features, in contrast to task irrelevant features. In contrast, top-down inhibition has been observed upon facing an expected stimulus, compared to an unexpected one (see [41] for a larger discussion about differences in attention and expectation).

This phenomenon of top-down inhibition, often referred to as “explaining away” in predictive coding literature, involves the modulation of lower-level cortical areas. This modulation serves to subtract neural activity patterns that align with prior brain information. Consequently, top-down inhibition leads to a reduction in information transfer when predictable inputs are received by the neuron.

This mechanism, characterized by a decrease in information transfer with increasing predictability of inputs, is manifested by the presence of negative LSTC. Our findings provide evidence supporting the existence of this inhibitory mechanism across cortical connections. Notably, detection was only feasible when computing LSTC at the resolution level of spiking activity. However, just a small fraction (33.33%) of negative LSTC in the spiking activity of the mouse visual system was detectable by our information-theoretic approach.

Our framework builds upon the concept of representation and error units, where error units compute prediction errors. These errors represent the residual neural activity after subtracting patterns consistent with prior brain information. The remaining neural activity, i.e., prediction errors, is then propagated up the cortical hierarchy, with the communication of these errors captured by negative LSTC.

An alternative neural implementation of explaining away, dendritic predictive coding [42], was proposed omitting the need for error units. The absence of these units would imply an absence of negative LSTC. Then, should evidence supporting dendritic predictive coding be reported, we suggest it could be complemented with a study of LSTC at the cellular level.

Moreover, our results not only support the presence of top-down inhibitory mechanisms but also a top-down enhancement with increasing predictability of neural inputs. This later case, captured by the positive LSTC observed both in LFP and spiking activity, aligns with theories such as ART or the biased competition model. These theories propose that cortical function relies on top-down excitation of neurons encoding stimulus features that are predictable within a given context.

Crucially, the two coding strategies described above are functionally equivalent [43]. They both rely on a predictive model instantiated in higher cortical areas, which encode the causes of sensory stimuli and modulate lower-level regions. However, our framework provides insights at the algorithmic level by representing neural processing computations in terms of information transfer and storage, thereby distinguishing between error and reliability coding strategies.

#### 4.3 The high density of cortical connections constrains the observability of information transfer from individual sources

In this study, we tested the presence of predictive processing coding strategies with an information-theoretic measure, namely the LSTC. This measure was computed for individual cortico-cortical connections, indicating the transfer of information between each pair of source and target brain areas. The density of brain connections, reported to be approximately 66% [44], imposes constraints on the amount of information transferred through each link — i.e., connections.

Specifically, this density was reported after examining in macaque monkeys the in-degree connectivity from 91 to 29 brain areas (Figure 1 in [44] depicts the corresponding areas), revealing 26 to 87 source areas linked to each target area (Figure 11 in [44]).

This high density of connections imposes limitations on the information transferred through individual links, with each link transferring only a small fraction of all cortical inputs to a given brain area. This limitation in information transfer may contribute to the observed small values — below r=0.10 — of the LSTC.

In addition, the small LSTC here presented are consistent with findings from a previous study [10] which reported values below r=0.10 for 9 out of 14 sub-cortical connections; and showed an increase in LSTC with the strength of connections (Figure 3 in [10]).

#### 4.4 A note on the absence of modulation of transfer entropy by external, macroscopic predictability

This study was explicitly aimed at testing predictive coding theories without recurrence to the experimenter’s views on what a certain brain area should predict, and to which violations of such prediction it should react. Nevertheless, the data taken for our iEEG analyses were taken from an experiment where external, macroscopic predictability of the stimuli was indeed explicitly manipulated. Thus, it is fair to ask whether these external manipulations resulted in differences in cortical information transfer as well, as predicted e.g. by the classic hierarchical ‘predicting away’-account of predictive coding. We tested for such differences using both a Bayesian logistic and linear regression (see S1 Text). For the logistic regression, we modelled the probability of the stimulus condition (i.e., expected or unexpected) as a function of TE and direction of connection (bottom-up, top-down); for the linear regression, we modelled the TE as a function of the condition and direction of connection. In these analysis we did not find evidence of relation between intracortical transfer entropy and stimulus predictability (supporting information S1 Fig and S4 Fig). While such a relation is clearly predicted by some popular theories of how predictive coding maybe implemented at the neural level [3], not all theories require macroscopic transmission of error signals (e.g. [42]). Thus, our results may provide first, albeit highly tentative evidence for the latter family of theories that do not require long-range information transfer to change in relation to the predictability of external or neural inputs.

However, even if classic accounts of predictive coding – where error signals are transmitted upwards in the cortical hierarchy – held, changes in information transfer in relation with changes of stimulus predictability may not be visible in our data for a number of reasons:

1. Due to constraints on the network size for which mTE could be computed we focused on sites which showed activity increases in response to changes in stimulus predictability (‘error sites’). It is possible that the increased information transfer which is predicted by explaining-away type of predictive coding theories does not happen predominantly *between these error sites*, but between error sites and other sites of the network.
2. Transfer entropy estimation from finite empirical data is known to be a difficult estimation problem, and it is also well known that establishing the presence of information transfer (detecting links) is much easier than getting reliable quantitative estimates of the amount of transferred information. Thus, our negative findings with respect to a modulation of information transfer by experimental manipulation may be due to a lack of statistical power (Large variance in TE estimates combined with only a few significant links entering the regression analysis as samples).
3. Furthermore, the brain areas recorded from may process information other than stimulus and stimulus-predictability information, with the latter only forming a small part of the total transferred information, also in this case the sensitivity of our analyses may be too limited to detect effects).

### 5 Conclusion

We tested the presence of predictive processing (PP) coding strategies at the cortical level in the human brain and mouse visual system using an information theoretic framework able to quantify two core PP concepts, i.e, information transfer and predictability. We quantify information transfer using Transfer Entropy and predictability using Active Information Storage. The correlation of these two measures, named Local Storage Transfer Correlations, was proposed as a signature of predictive processing given that it naturally captures the enhancement (positive correlation) or suppression (negative correlation) of information transfer with neural-predictability. In this way, the sign of these correlations allows teasing apart two types of predictive processing strategies: error-coding, with a negative correlation, and reliability-coding, with a positive correlation. We observed a reliability-coding strategy unfolded in 17 cortico-cortical connections detected using transfer entropy in the human intracranial recordings; these positive correlations were present in frontal, parietal and temporal brain areas. In contrast, the mouse visual system presented 9 cortical connections which implement both a reliability-coding (66.67% of the connections) and error-coding (33.33% of the connections) strategy.

## 6 Acknowledgments

We would like to thank the technologists team of the Epilepsy Monitoring Unit at the University Medical Center Göttingen and Manuel Hewitt for the ongoing and important support.

