## Supplementary Text and Figures for "Correlations of neural predictability and information transfer in cortex and their relation to predictive coding"

### Supporting information

Our information-theoretic framework is suitable for studying predictive processing by estimating both information transfer and predictability from neural activity during recordings with uncontrolled stimulus predictability. However, in this work, we analyzed iEEG recordings from human epileptic patients during a face-processing task, where the stimulus predictability was controlled. This allowed us to study how stimulus predictability modulates transfer entropy intensity (i.e., the amount of information transfer).

We implemented two Bayesian regressions: logistic and linear. For the logistic regression, we estimated the probability of the stimulus being expected based on transfer entropy intensity (TEi) and the direction of connection. For the linear regression, we aimed to predict transfer entropy intensity from the experimental condition (i.e., whether the stimulus was expected or unexpected) and the direction of connection.

#### Logistic regression: Transfer entropy intensity and direction of connection as predictors of stimulus predictability

We used a logistic regression model to estimate the probability of the stimulus being expected based on TEi and the direction of connection. Our model used as input data the connections detected via transfer entropy network analysis on iEEG recordings from human epileptic patients. We applied logistic regression to predict the condition – i.e., whether the stimulus was expected or unexpected.

For this, we used the Bernoulli distribution as the likelihood function, with the probability p of the stimulus being expected described as a function of both TEi and the direction of connection (see Eq. S1). The direction of connection captures the influence of the cortical hierarchy on the probability of the stimulus being expected. Represented as a categorical variable, the direction of connection was coded as either 1 (top-down) or -1 (bottom-up). Connections were classified as top-down if the source brain area was at a lower level in the cortical hierarchy than the target brain area, whereas bottom-up connections referred to the opposite direction.

Their classification (see Table S2) was based solely on the literature [1, 2, 3], allowing us to classify 9 out of 17 connections, which constituted the data for our models. As a control, we also fit two additional models: one with only transfer entropy intensity as a predictor (transfer entropy model) and another without any predictors (null model).

$$p = \frac{1}{1 + e^{-(\beta_0 + \beta_{TEi} \cdot TEi + \beta_d \cdot d + \beta_{interaction} \cdot TEi \cdot d)}} \qquad (1)$$

where $\beta_0$ is an intercept, $\beta_{TEi}$ a parameter capturing the influence of TEi, $\beta_d$ a parameter capturing the influence of direction of connection, $\beta_{interaction}$ a parameter capturing the influence of the interaction between TEi and direction. See Table S1 for the priors.

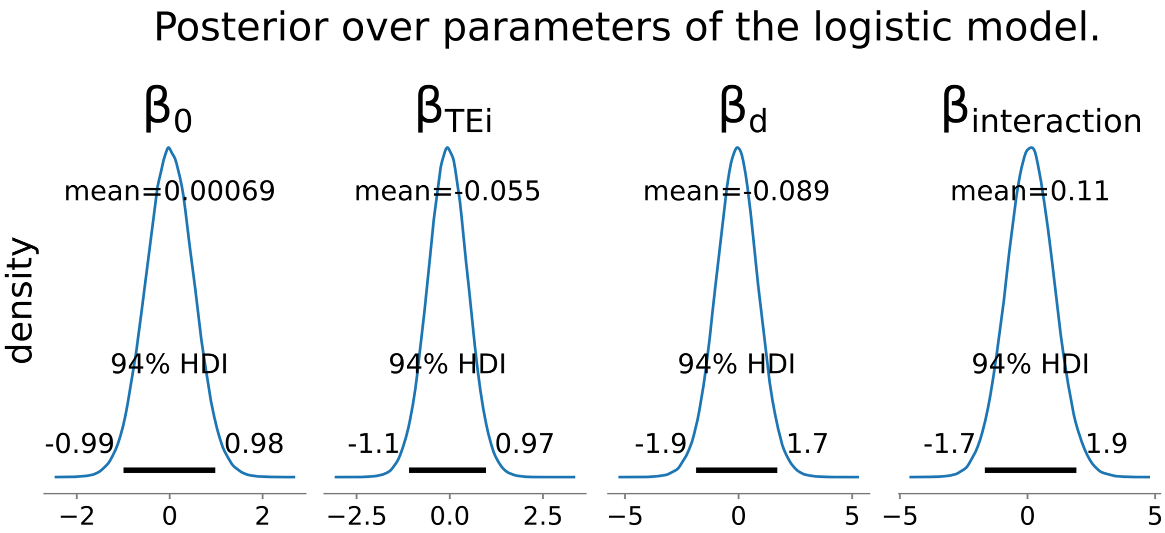


**Figure 1.** Posterior over parameters of logistic model. The condition (i.e., expected or unexpected) was modelled using a Bernoulli likelihood with the probability $p$ of stimulus being expected modelled as a function of intensity of transfer entropy (with parameter $\beta_{TEi}$), direction of connection (with parameter $\beta_d$) and an interaction term between these two (with parameter $\beta_{interaction}$). For each posterior, the 94% highest density interval (HDI) was included.

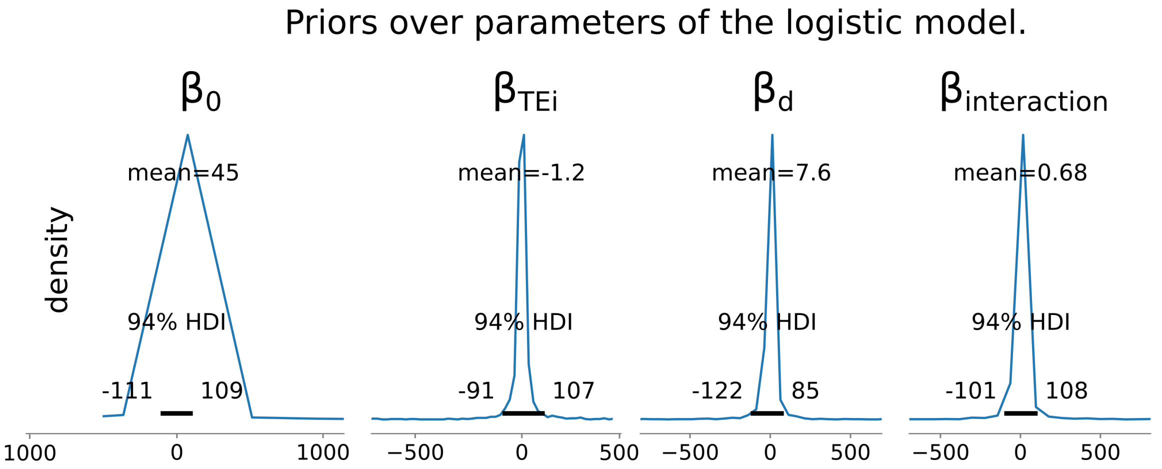


**Figure 2.** Priors over parameters of logistic model. See Table (S1) for the definition of priors' distributions. The parameter $\beta_{TEi}$ captures the influence of the intensity of transfer entropy, $\beta_d$ that of the direction of connection and $\beta_{interaction}$ an interaction term between these two predictors. For each prior, the 94% highest density interval (HDI) was included.

**Absence of prediction of stimulus predictability by transfer entropy and direction of connection.**

Our evidence shows that the probability of the stimulus being expected is independent of both TEi ([-1.09,0.97], 94% CI) and direction of connection ([-1.87,1.70], 94& CI). Furthermore, the model with only an intercept (i.e., the null model) was the best, as indicated by the leave-out-out (LOO) cross-validation-based Bayesian model comparison [4] (Fig S3). We observed that the null model outperformed both the *transfer entropy & direction* model and the *transfer entropy* model. Additionally, a comparison between the posteriors (Fig S1) and priors (Fig S2) of the parameters shows that, although the 94% highest density interval was reduced (e.g., for $\beta_{TEi}$ from [-91, 107] to [-1.10, 0.97]) the posteriors remained symmetric around zero. This indicates an absence of data influence in shifting the posterior toward either positive or negative values.

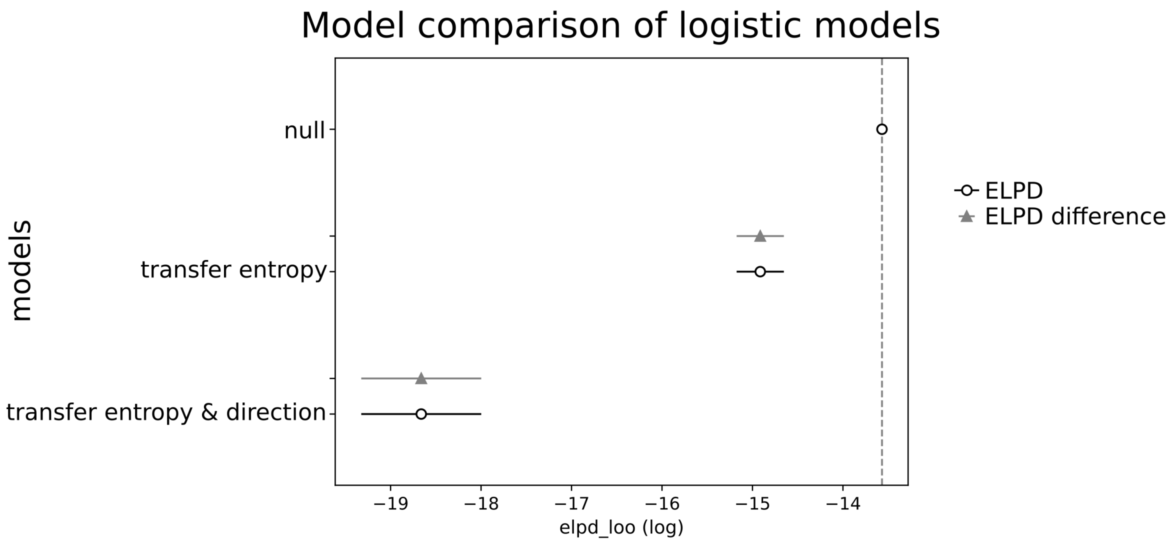


**Figure 3.** Model comparison between three logistic models: *transfer entropy & direction* model, *transfer entropy* model, *null* model. The *transfer entropy & direction* model expressed in Eq S1 had both the transfer entropy intensity and the direction of connection as predictors of the probability of stimulus being expected. The *transfer entropy* only included the transfer entropy intensity as predictor and the *null* model just an intercept. Models were ranked based on their expected log pointwise predictive density (ELPD) estimated using the leave-out-out cross-validation (LOO) method. Grey triangles represent the difference between the ELPD of a given model and the best one. We observed that the best model (i.e., lowest elpd loo) was the null model supporting the absence of influence of both TEi and direction of connection on predicting the condition.

### Linear regression: Stimulus predictability and connection's direction as predictors of transfer entropy intensity

For the linear regression, we used TEi as the data, modeling it with a Normal distribution as the likelihood. We modeled the mean, $\mu$, of the Normal distribution using both the condition (i.e., stimulus expected or unexpected) and the direction of connection as predictors (see Eq S2). In both cases, we used categorical variables with the same representation as in the logistic models: for the condition, 1 (expected) and 0 (unexpected); and for the direction of connection, 1 (top-down) and -1 (bottom-up). As a control, we also fitted a model with only the condition as predictors and a null model without any predictors.

$$\mu_{TEi} = \beta_0 + \beta_c \cdot c + \beta_d \cdot d + \beta_{interaction} \cdot c \cdot d \quad (2)$$

where $\beta_0$ is an intercept, $\beta_c$ a parameter associated to the condition, $\beta_d$ a parameter associated to the direction of connection, $\beta_{interaction}$ a parameter associated to the interaction between the condition and direction of connection. For the priors used for each parameter see Table S5.

#### Absence of modulation in transfer entropy intensity by stimulus predictability and direction of connection.

Our evidence shows that TEi is independent of both the condition ([-0.61, 0.54], 94% CI) and the direction of connection ([-0.57, 0.56], 94% CI).

As with the logistic regression, the model with only an intercept (i.e., the null model) was the best, as indicated by the leave-out-out (LOO) cross-validation-based Bayesian model comparison [4] (Fig S6). A similar pattern was observed as in the logistic regression, where the null model outperformed both the *condition & direction* model and the *condition* model.

| Parameter | Prior distribution |
|---|---|
| $\beta_0$ | Cauchy($\alpha$=0, $\beta$=10) |
| $\beta_{TEi}$ | Cauchy($\alpha$=0, $\beta$=10) |
| $\beta_d$ | Cauchy($\alpha$=0, $\beta$=10) |
| $\beta_{interaction}$ | Cauchy($\alpha$=0, $\beta$=10) |

**Table 1.** Priors for the parameters of the logistic regression. We chose wide priors to prevent prior beliefs from overly influencing the posteriors given the limited amount of data we had. Data was z-score before modeling. The functional form of the Cauchy distribution is $f(x \mid \alpha, \beta) = \frac{1}{\pi\beta[1+(\frac{x-\alpha}{\beta})^2]}$.

| Patient | Source | Target | Direction of connection |
|---|---|---|---|
| 2418517735 | L Rostral Middle Frontal | L Inferior Temporal | Top-Down [1] |
| 3506483969 | L Hippocampus | L Amygdala | Top-Down |
| 3506483969 | L Hippocampus | L Parahippocampal | Top-Down [2] |
| 5368380838 | R Caudal Middle Frontal | R Hippocampus | Bottom-Up |
| 5368380838 | R Supramarginal | R Insula | Bottom-Up [3] |
| 5368380838 | R Rostral Middle Frontal | R Supramarginal | Top-Down [3] |
| 5921933672 | L Fusiform | L Middle Temporal | Top-Down |
| 5921933672 | L Middle Temporal | L Fusiform | Bottom-Up |
| 9617561518 | L Inferior Parietal | L Insula | Bottom-Up [3] |

**Table 2.** Classification of links as Top-Down or Bottom-Up based on the levels in the cortical hierarchy of the connected brain areas. Top-Down links when the source brain area is in a higher level than the target brain area. Bottom-up if the source brain area is in a lower level than the target brain area.

| Parameter | Prior distribution |
|---|---|
| $\sigma$ | HalfCauchy($\beta$=100) |
| $\beta_0$ | Normal($\mu$=0, $\sigma$=100) |
| $\beta_c$ | Normal($\mu$=0, $\sigma$=100) |
| $\beta_d$ | Normal($\mu$=0, $\sigma$=100) |
| $\beta_{interaction}$ | Normal($\mu$=0, $\sigma$=100) |

**Table 3.** Priors for the parameters of the linear regression. We chose wide priors to prevent prior beliefs from overly influencing the posteriors given the limited amount of data we had. Data was z-score before modeling.

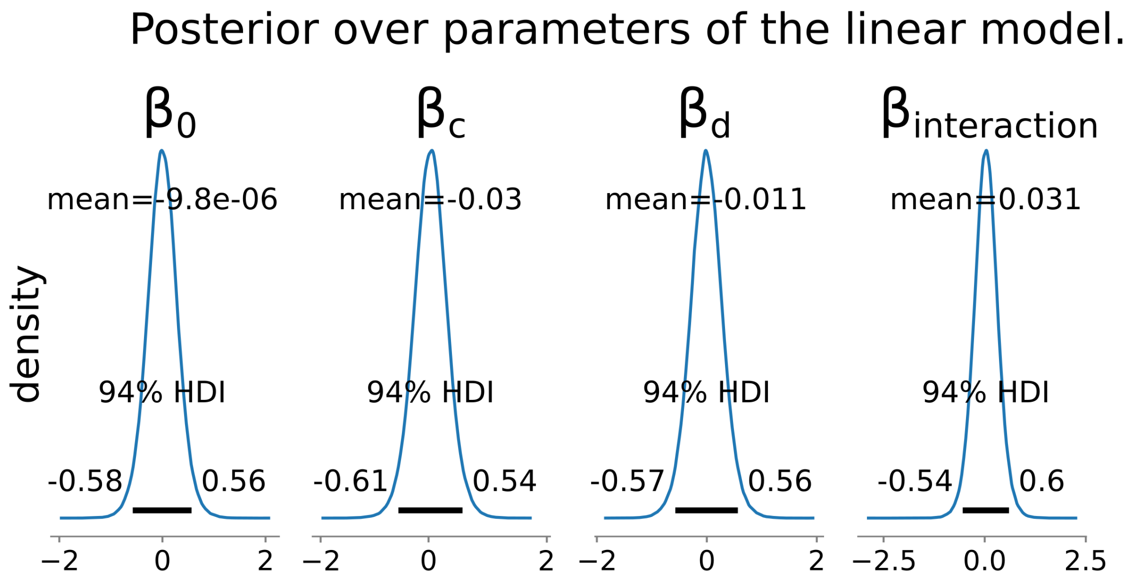


**Figure 4.** Posterior over parameters of linear model. The transfer entropy intensity was modelled using a Normal likelihood with its mean

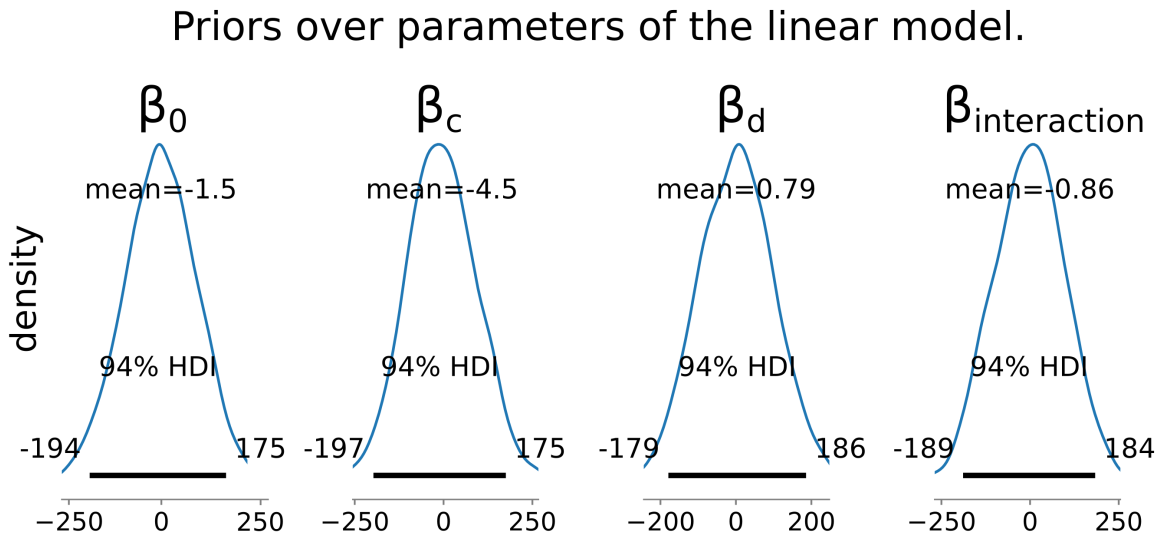


**Figure 5.** Priors over parameters of linear model. See Table (S5) for the definition of priors' distributions. The parameter $\beta_c$ captures the influence of the condition, $\beta_d$ that of the direction of connection and $\beta_{interaction}$ an interaction term between these two predictors. For each prior, the 94% highest density interval (HDI) was included.

An analysis of the posteriors and priors (Fig S4 and S5) leads to the same conclusions as for the logistic regression – i.e., the highest density intervals were reduced but remained symmetric around zero.

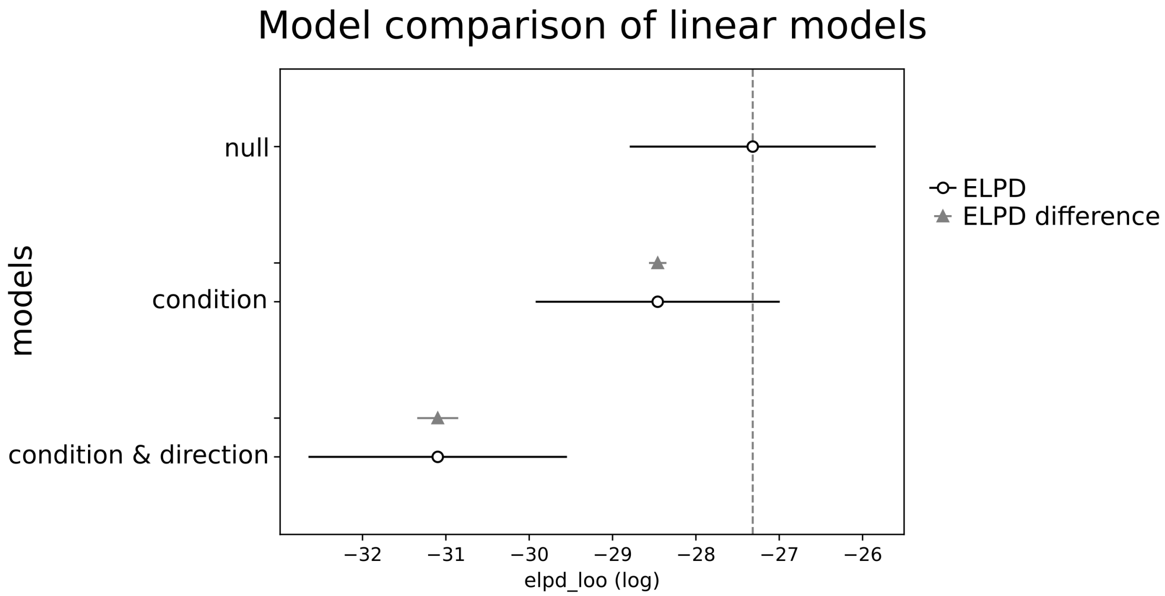


**Figure 6.** Model comparison between three linear models: *condition & direction* model, *condition* model, *null* model. The *condition & direction* model expressed in Eq S2 had both the condition and the direction of connection as predictors of the transfer entropy intensity. The *condition* only included the condition as predictor and the *null* model just an intercept. Models were ranked based on their expected log pointwise predictive density (ELPD) estimated using the leave-out-out cross-validation (LOO) method. Grey triangles represent the difference between the ELPD of a given model and the best one. We observed that the best model (i.e., lowest elpd loo) was the null model supporting the absence of influence of both the condition and direction of connection on predicting the transfer entropy intensity.
